# Negative Feedback Facilitates Temperature Robustness in Biomolecular Circuit Dynamics

**DOI:** 10.1101/007385

**Authors:** Shaunak Sen, Richard M. Murray

## Abstract

Temporal dynamics in many biomolecular circuits can change with temperature because of the temperature dependence of underlying reaction rate parameters. It is generally unclear what circuit mechanisms can inherently facilitate robustness in the dynamics to variations in temperature. Here, we address this issue using a combination of mathematical models and experimental measurements in a cell-free transcription-translation system. We find that negative transcriptional feedback can reduce the effect of temperature variation on circuit dynamics. Further, we find that effective negative feedback due to first-order degradation mechanisms can also enable such a temperature robustness effect. Finally, we estimate temperature dependence of key parameters mediating such negative feedback mechanisms. These results should be useful in the design of temperature robust circuit dynamics.

## 1 Introduction

Understanding and designing robust performance in circuit function has been an important goal ever since the initial demonstrations of synthetic circuits that demonstrated the ability to engineer complex dynamical behavior [5,6]. Efforts to achieve this goal of functional robustness have proceeded at multiple levels. At one level, there have been efforts towards a systematic characterization of circuit elements so that the uncertainty in underlying parameters is minimized prior to their use in circuit construction (for example, [8, 9]). At another level, insulator elements have been proposed to shield the function of individual components when they are connected into larger circuits [4]. Indeed, designing biomolecular circuits of the size and functional robustness on par with naturally occurring circuits can be viewed as the holy grail of synthetic biology.

In particular, in the study of naturally occurring biomolecular circuits, robustness to the important environmental variable of temperature has received much attention. This robustness may be desired for a dynamic phenotype as well as for an equilibrium phenotype. For example, a classical problem is how the dynamic phenotype corresponding to the period of biomolecular oscillators is almost independent of temperature [10]. Similarly, equilibrium phenotypes such as the adaptation level in bacterial chemotaxis can also be robust to changes in temperature [11]. In both these cases, the opposing effect of rates with similar temperature dependence is believed to be the key mechanism underlying temperature robustness. Indeed, robustness to temperature is likely to be similarly important for biomolecular circuit design. Recent work has characterized the temperature dependence in biomolecular circuits [13] as well as designed circuits with temperature robust phenotypes [7]. In general, for biomolecular circuit design, a key challenge is to identify mechanisms that intrinsically suppress variations due to temperature.

A key mechanism that enables robustness in multiple dynamic contexts is that of negative feedback. Negative feedback is frequently used in control engineering design to ensure that the output of a system stays at its desired value despite disturbances, static or dynamic, that act on the system [1]. Analogous uses of negative feedback have been reported in investigations of biomolecular circuits. Experiments with simple circuit designs have been used to show that negative feedback can reduce the effect of noise in the equilibrium value of concentrations [3]. Further, negative feedback is also operational in biomolecular signaling systems with adaptation, wherein the output returns to a fixed value after a transient response to a change in input [2, 15]. Given these, a natural question to ask is how negative feedback in a biomolecular circuit responds to a variation in temperature.

Here we ask whether negative feedback can facilitate robustness to temperature in biomolecular circuit dynamics. To address this, we use a combination of mathematical modeling and experimental measurements in a cell-free transcription-translation system. We find that negative feedback dynamics can suppress variation in circuit dynamics due to a change in temperature. Next, we measure circuit dynamics at different temperatures and find that they support this finding. Finally, we use this data to estimate the temperature dependence of key circuit parameters. These results should help in design of systems with temperature robustness properties.

## 2 Results

### 2.1 Mathematical Modeling

Consider a simple model of a negative transcriptional feedback circuit, where a protein *X* is a transcription factor that can repress its own transcription,

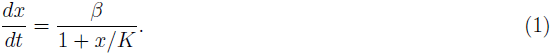

Here, *x* denotes the concentration of the protein, *β* denotes the maximal production rate, and *K* denotes the dissociation constant for the binding of *x* to its own promoter. As the concentration of *X* increases from a zero initial condition, the rate of production transitions from a constant *β* (when *x* < *K*) to an *x*-dependent rate *βK/x*. In this state, when the negative feedback becomes operational, the equation simplifies to,

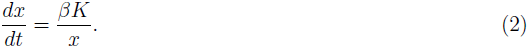

Using this simplification, the time evolution of *x* can be approximated as,

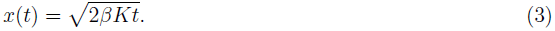

The nonlinear dependence of *x* on *t* arises due to the negative feedback mechanism. This is in contrast to the linear dependence when *x* is expressed from a constitutive promoter, whose promoter activity is constant with time. To see this, consider the equation,

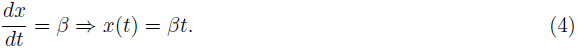

Using these expressions, we investigate the effect of a small variation in temperature, *T→T+*Δ*T*, on the temporal dynamics. Corresponding to this variation, the change in trajectory is, 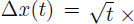 (difference in parameters). We note that the variation in trajectory scales only as the square root of time. In contrast, for the case of a constitutive promoter, the difference scales with time, Δ*x*(*t*) = *t* × (difference in parameters). Therefore, we note that this negative feedback mechanism can facilitate robustness to a variation in temperature by means of a smaller scaling of the corresponding difference in the trajectory with time. This effect is solely dependent on the nonlinearity introduced by the mechanism and is apparent in the circuit dynamics.

To check whether this holds in the complete model, we solve Eqn. (1) exactly,

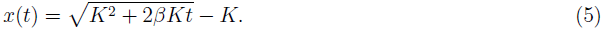

Initially (*t < K/β*), the concentration of *X* increases linearly with time, *x*(*t*) = *βt*. Later (*t > K/β*), the concentration of *X* has a square root dependence on time, 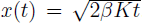. This corresponds to the case where the negative feedback mechanism becomes operational. Therefore, in this regime of the general model, the temperature robustness effect comes into play. In fact, the stronger the binding of *X* to its own promoter, i.e. the smaller the value of *K*, earlier is the transition of *x* to a square root dependence on time. This means that, the stronger the negative feedback, the earlier is the onset of this temperature robustness effect.

### 2.2 Experimental Measurements

In order to check for signatures of the above discussed effect, we performed experimental measurements. These experiments were performed in a cell-free transcription translation system, which offers a conveniently fast platform to analyze biomolecular circuit dynamics in cell-like contexts [12, 14].

The negative transcriptional feedback circuit used for the experiment contained a transcriptional repressor protein TetR expressed under a self-repressible promoter (Fig. 1). In this circuit, TetR was fused to a green fluorescent protein variant deGFP to allow for measurement of circuit dynamics, using excitation and emission at wavelengths 485 nm and 525 nm, respectively. These measurements were performed in a multilabel platereader (BioTek Synergy H1) with measurement intervals set at 3 minutes for a total duration of 10 hours. These were then repeated at the different desired temperatures. Further, using the inducer anhydrotetracycline (aTc), an inhibitor of TetR’s ability to bind DNA, two different negative feedback strengths were measured. The aTc concentration of 0.5 *µ*g/ml corresponded to relatively stronger negative feedback and that of 5 *µ*g/ml corresponded to relatively weaker negative feedback strength. These experimental data are shown in Fig. 1. Different colors represent different temperatures, which, based on the measurement logs, had mean values *T*_1_ = 26*°C* (Fig. 1, blue), *T*_2_ = 29*°C* (Fig. 1, cyan), *T*_3_ = 33*°C* (Fig. 1, magenta), *T*_4_ = 37*°C* (Fig. 1, red).

**Figure 1:**
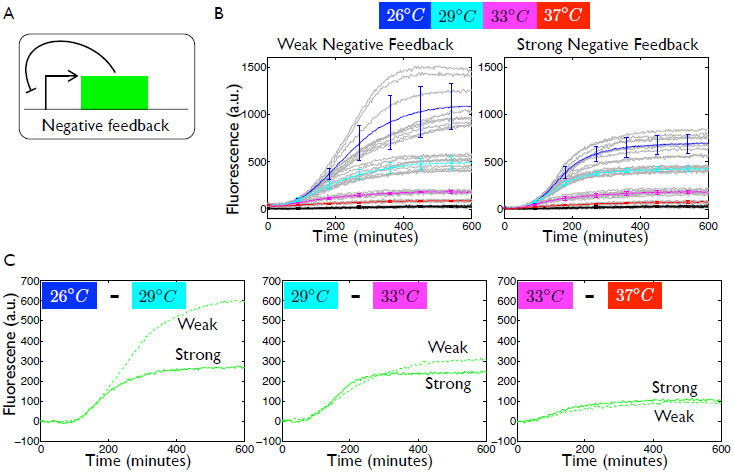
Temperature robustness in the temporal dynamics of a transcriptional negative feedback circuit. A. Schematic of the negative feedback circuit. B. Grey lines show the raw data that are acquired for each temperature, in triplicate on each day, and on three separate days. Colored lines with errorbars superimposed on the raw data are mean and standard deviation, with these statistics plotted at 30 minute intervals. The colors blue, cyan, magenta, and red represent the mean values of measured temperatures *T*_1_ = 26*°C*, *T*_2_ = 29*°C*, *T*_3_ = 33*°C*, and *T*_4_ = 37*°C*, respectively. Black traces represent the background of a reaction with no DNA. C. Panels represent differences between the mean traces for indicated temperatures. Solid and dashed green lines represent the differences for the strong and weak negative feedback cases, respectively.

To estimate the variation in the dynamic trajectory due to a variation in temperature, we computed the differences in the mean values of the above data over time. From modeling considerations presented above, we expect the reduced scaling with time to appear earlier for a stronger negative feedback. This differences were computed between the mean values acquired at *T*_1_ = 25*^o^*C and *T*_2_ = 29*^o^*C, at *T*_2_ = 29*^o^*C and *T*_3_ = 33*^o^*C, and at *T*_3_ = 33*^o^*C and *T*_4_ = 37*^o^*C (Fig. 1C). Furthermore, these were computed for the strong negative feedback case as well as the weak negative feedback case. For the first two differences, we find that the scaling with time reduces earlier for the strong negative feedback case compared to the weaker negative feedback case, consistent with what is expected from modeling considerations. For the last difference, it is difficult to differentiate between the scalings owing to the low signal amplitude. Based on this analysis, we conclude that there is experimental support for an inherent temperature robustness property due to transcriptional negative feedback dynamics.

**Effect of saturation.** We noted that traces in the above data plateau as time increases. This saturation feature is not present in the simple models considered above. Here, we investigate the effect of this saturation on the temperature robustness property. In fact, the same saturation effect is also present in the dynamics of a constitutive promoter expressing deGFP (Fig. 2). This implies that the saturation feature is an inherent consequence of the cell-free transcription-translation system. One possibility for this is the decay of resources in the expression system. To model this, we augmented the model with a first-order decay of the expression machinery as follows,

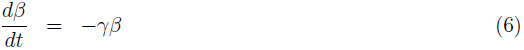

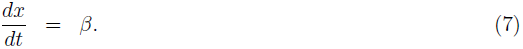

Here, *β* is a lumped parameter for the resource that decays over time. The solution 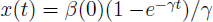 increases linearly with time initially (*t <* 1*/γ*) and then saturates to *β*(0)*/γ* for large time (*t >* 1*/γ*). Therefore, the solution exhibits a saturation effect for the constitutive promoter.

**Figure 2:**
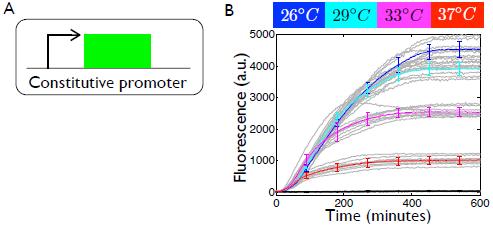
Saturation effect in the constitutive promoter dynamics. A. Schematic of the constitutive promoter circuit. B. Grey lines show the raw data that are acquired for each temperature, in triplicate on each day, and on three separate days. Colored lines with errorbars superimposed on the raw data are mean and standard deviation, with these statistics plotted at 30 minute intervals. Black traces represent the background of a reaction with no DNA.

To see if a similar resource decay in the negative feedback model can generate robustness to variation in temperature similar to that see above, we considered the augmented model,

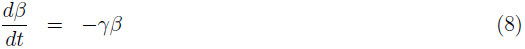

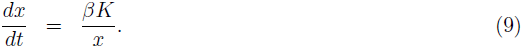

The solution is 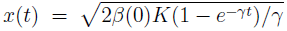. It can be seen that initially 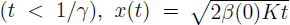, which has the same reduced scaling with time as seen above.Therefore, the same temperature robustness effect can be seen even when a resource decay is explicitly augmented in the model.

**Effective negative feedback due to resource degradation.** Based on analysis of the above Eqns. (6)-(9), we note that the presence of the degradation dynamics results in a sub-linear dependence of the trajectory on time. This arises due to the presence of the 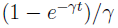 term in the solutions. Indeed the solution for the constitutive promoter with resource degradation is 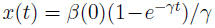. This trajectory initially increases linearly with time and then eventually saturates. Between these two limits, the dependence of the trajectory on time is sub-linear. Mathematically, these degradation dynamics can be viewed as arising from an implicit negative feedback in the model, where the net degradation is directly proportional to the amount of resource. Increasing the amount of effective negative feedback through an increase in *γ* results in an earlier onset of the sub-linear dependence on time, which should reduce the variation in traces as a function of temperature. Therefore, the effective negative feedback due to degradation dynamics can have a similar effect as transcriptional negative feedback in reducing the effect of transient response of a variation in temperature.

### 2.3 Temperature Dependence of Parameters

In the above results, we investigated how robustness in the trajectory to a variation in temperature is facilitated by inherent features in the mechanism. In general, the circuit parameters themselves can change with temperature. The experimental measurements shown above can be used to estimate the temperature dependence of key circuit parameters. Next, we used basic parametric fits to obtain these temperature dependencies for the constitutive promoter as well as the transcriptional negative feedback circuit dynamics.

First, for the constitutive promoter, we aimed to estimate the lumped parameter *β* corresponding to the production rate in Eqn. (4). A simple way to estimate this is through the slope of the deGFP trajectory. We use a sliding window of width 30 minutes to calculate the slope at each time point and estimate *β* as the maximum slope over the whole time duration. This provides an estimate of *β* in terms of the maximum production rate. Deviations from the maxima are due to non-idealities in the system, for example, the saturation of traces, possibly due to resource decay, leads to an eventual decline of the slope from this maximum value. We perform this parameter estimation for the mean fit as well as for each of the individual traces at different temperatures (Fig. 3). We find that the estimated production strength is approximately constant in the range 25–33*^o^* C and then halves at 37*^o^* C. These results provide an estimate of how the strength of a constitutive promoter changes with temperature.

**Figure 3:**
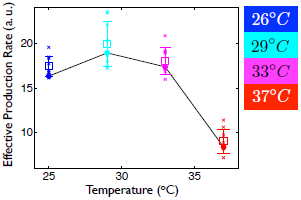
Estimate of the temperature dependence of the effective production strength in a constitutive promoter. Circles represent the estimates obtained from the mean traces. Crosses represent the estimates obtained from the individual traces. Open squares with error bars represent the mean and standard deviation of the crosses. The colors blue, cyan, magenta, and red represent the mean values of measured temperatures *T*_1_ = 26*°C*, *T*_2_ = 29*°C*, *T*_3_ = 33*°C*, and *T*_4_ = 37*°C*, respectively.

Next, we use the traces obtained for negative feedback to estimate temperature dependence of respective lumped parameters in Eqn. (1). As above, we estimate effective production strength *β* from the maximum slope of these traces. To estimate effective binding constant *K*, we rearrange the functional form of the solution 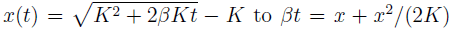, where *β* has been estimated from the maximum slope. This is then fit using a curve-fit from the time that the trace reaches maximum. As above, this is done for each of the individual traces as well as for the mean trace. We report these two fits for both parameters and for both negative feedback scenarios in Fig.4. These fits show that both maximal promoter activities are similar and decrease with temperature, approximately halving every 4*^o^*C. It is interesting to note that the estimate of the production strength of the constitutive promoter is different from this. Similarly, the estimates of binding constants also decay with temperature. As expected, the binding constant in the stronger negative feedback scenario is typically smaller than the one with weaker negative feedback. These result provide estimates of the temperature dependence of key lumped parameters in the negative feedback circuit.

**Figure 4:**
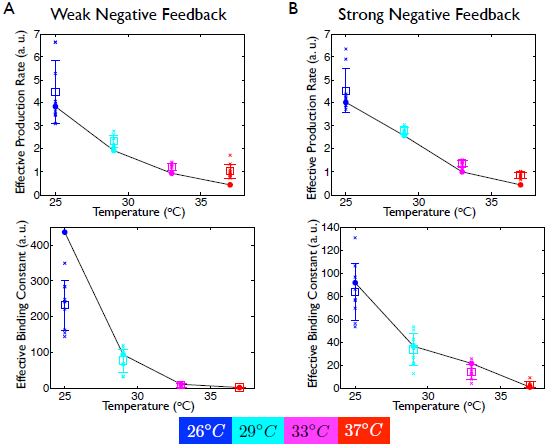
Estimate of the temperature dependence of the effective production rate and effective binding constant in a negative feedback circuit for (A) Weak negative feedback and (B) Strong negative feedback. Circles represent the estimates obtained from the mean traces. Crosses represent the estimates obtained from the individual traces. Open squares with error bars represent the mean and standard deviation of the crosses.

## 3 Discussion

Investigating mechanisms that can inherently facilitate functional robustness against variations in temperature is an important problem for biomolecular circuit design. Using a combination of simple mathematical modeling and measurements of circuit dynamics in a cell-free transcription-translation system, we present three key results. First, we identify temporal dynamics due to negative transcriptional feedback as those that can inherently suppress variation due to temperature. Second, we present measurements of these circuit dynamics for different temperatures in support of the above finding. Third, we use our experimental data to get an estimate of temperature dependence of key parameters of the negative transcriptional feedback circuit. These results illustrate how robustness to temperature in dynamics can be facilitated in a commonly used biomolecular circuit module.

Robustness to a dynamic phenotype such as the change in trajectory over time is an interesting aspect emphasized in this study. Indeed, most studies on robustness focus on an equilibrium phenotype such as the activity level of a protein [2]. Another example where such a dynamic property arises is in the investigation of temperature compensation of circadian rhythms, where the period of oscillation is found to be robust to changes in temperature. Robustness to such dynamic properties are also of significance to the study of natural circuits and design of synthetic ones.

A challenging task for future work is to similarly investigate the effect of variation of temperature on other biomolecular circuit mechanisms. In particular, positive feedback mechanisms offer a natural class of systems for this purpose. Preliminary calculations suggest an opposite effect in their case, with a variation in temperature that is amplified in the dynamics. Together, these investigations will help shed further insight in the analysis of biomolecular oscillator dynamics, which frequently contain both positive and negative feedback loops. In fact, the cancellation of the opposite effects of a temperature variation on the dynamics of positive and negative feedback loops may contribute to improving our understanding of temperature compensation in the oscillator dynamics.

Ensuring robustness in biomolecular circuits is an important step in their design. Identifying and harnessing inherent robustness-enabling biomolecular mechanisms is one way to ensure this. Here, we have identified negative feedback as providing robustness of the trajectory to variations in the important environmental variable of temperature. These results should help in designing robustness to temperature in dynamic contexts as well as in understanding mechanisms underlying temperature robustness in naturally occurring biomolecular contexts.

## Acknowledgements

We gratefully acknowledge C. Hayes for her help with the experimental part of this work, especially for providing the constructs for the negative feedback circuit and the constitutive promoter as well as for the measurement protocol.

